# miRNome profiling of clonal stem cells in Ph^+^ CML

**DOI:** 10.1101/2020.03.16.989194

**Authors:** María Sol Ruiz, María Belén Sánchez, Simone Bonecker, Carolina Furtado, Daniel Koile, Patricio Yankilevich, Santiago Cranco, María del Rosario Custidiano, Josefina Freitas, Beatriz Moiraghi, Mariel Ana Peréz, Carolina Pavlovsky, Ana Inés Varela, Verónica Ventriglia, Julio César Sánchez Ávalos, Irene Larripa, Ilana Zalcberg, José Mordoh, Peter Valent, Michele Bianchini

## Abstract

Chronic myeloid leukemia (CML) is a myeloid stem cell neoplasm characterized by an expansion of myeloid progenitor cells and the presence of BCR-ABL1 oncoprotein. Since the introduction of specific BCR-ABL1 tyrosine kinase inhibitors (TKI), overall survival has improved significantly. However, under long-term therapy patients may have residual disease that originates from TKI-resistant leukemic stem cells (LSC). In this work, we analyzed the miRNome of CML LSC, normal hematopoietic stem cells (HSC) obtained from the same CML patients, and stem and progenitor cells obtained from healthy donors (HD) by next-generation sequencing. We detected a global decrease of microRNA levels in LSC and HSC from CML patients, and decreased levels of microRNAs and snoRNAs from a genomic cluster in chromosome 14, suggesting a mechanism of silencing of multiple non-coding RNAs. Surprisingly, HSC from CML patients, despite the absence of *BCR-ABL1* expression, showed an altered miRNome. *In silico* analysis revealed an association between validated microRNAs and multiple metabolic pathways, suggesting that these molecules may be mediators of the previously reported dysregulation of LSC metabolism. This is the first report of the LSC miRNome that distinguishes between *BCR-ABL1*^*+*^ LSC and their *BCR-ABL1*^*-*^ counterparts, providing valuable data for future studies.

## Introduction

Chronic myeloid leukemia (CML) originates from a hematopoietic stem cell (HSC) that acquires the reciprocal translocation t(9;22) (q34;q11) and thus the Philadelphia chromosome (Ph)^1^. The resulting fusion gene, *BCR-ABL1*, encodes an oncogenic protein with constitutive tyrosine kinase activity. Treatment of CML patients was revolutionized by the introduction of specific tyrosine kinase inhibitors (TKI), like imatinib, nilotinib or dasatinib. These TKI effectively induce apoptosis in leukemic cells in patients with CML^2^. However, the response of patients to TKI treatment is heterogeneous, and about 40% of imatinib-treated patients require a switch of TKI due to intolerance or resistance to treatment^3^. Other patients with optimal response to TKI show persistence of the leukemic clone, even after several years of treatment^4^. A subset of TKI-treated CML patients can achieve a deep molecular response during therapy^3^. However, only half of them or even less can sustain a treatment-free remission^5–7^.

Leukemic stem cells (LSC) are defined as a population of cells that gives rise and maintains the leukemic clone^8^. The classical view of CML considers that LSC derive from the acquisition of BCR-ABL1 in a HSC^9,10^. However, BCR-ABL1 alone is unable to induce a leukemia^10,11^. Rather, additional molecular lesions and hits are required for full transformation of clonal pre-leukemic (stem) cells into fully malignant leukemic (stem) cells. Correspondingly, single-cell gene expression analysis revealed great heterogeneity within LSC populations^10,12^. Their normal counterparts, HSC, also constitute a heterogeneous population, and individual HSC exhibit certain properties related to their stem cell nature: self-renewal, quiescence, repopulation capacity, and differentiation potential^13,14^. The mechanisms underlying the regulation of such properties are not completely understood; however, they depend on both intrinsic (such as the levels of specific transcription factors) and extrinsic (such as signals coming from the bone marrow niche) factors^14,15^. In CML, most LSC and their subclones may be sensitive to TKI therapy. However, certain stem cell classes, especially pre-leukemic neoplastic stem cells may be resistant because they are slowly cycling cells and exhibit multiple forms of stem cell resistance^10,13^. Sometimes even LSC may survive TKI therapy and thus persist in CML patients. The persistence of LSC in patients under TKI therapy has fueled intensive research on this topic, in order to identify novel therapeutic targets that enable the complete eradication of the leukemic clone in all patients^3^. On the other hand, it is not clear whether the heterogeneity observed in the LSC population is related to different responses to TKI treatment.

In CML, recent reports have characterized the transcriptome of LSC^16,17^, protein networks of precursor cells (CD34^+^)^18^, and the metabolome of LSC^19^. Gene expression profiling of the primitive fraction of leukemic cells in CML patients revealed a transcriptional profile resembling normal CD34^+^ myeloid progenitor cells, with decreased levels of transcription factors involved in maintenance of stem-cell fate, suggesting loss of quiescence^17^. Single-cell RNA sequencing revealed an enrichment of gene sets related to mechanistic target of rapamycin kinase (*MTOR*), targets of E2F transcription factors, G2/M checkpoints, oxidative phosphorylation, and glycolysis-associated gene expression in *BCR-ABL1*^*+*^ stem cells^16^. However, little is known about microRNA-mediated regulation of gene expression in this population. MicroRNAs are small (19-25nt), non-coding RNAs that can regulate multiple targets, mainly by mRNA destabilization or inhibition of protein translation. They are evolutionary conserved, and have shown to be relevant for multiple physiological and pathological processes^20^. One report has shown the involvement of microRNAs in TKI sensitivity in CML LSC^21^. Recent advances in the characterization of aberrant expression of surface markers have allowed the prospective isolation of LSC and HSC from CML patients^12,22^. In this work, we aimed to characterize the miRNome of LSC and HSC isolated by fluorescence-activated cell sorting (FACS) from CML patients at diagnosis by small RNA-Next-Generation Sequencing (NGS), in order to identify differential molecular mechanisms that contribute to unravel LSC biology, and the possible therapeutic implications of such differences. We observed a global decrease in microRNA levels in LSC and putative HSC from CML patients in comparison with HSC obtained from healthy donors (HD). Surprisingly, compared to HSC from HD, we detected decreased levels in LSC of microRNAs and snoRNAs belonging to a genomic cluster located in chromosome 14 (14q.32) that contains imprinted genes, suggesting an epigenetic mechanism of silencing of multiple non-coding RNAs. Finally, we validated a group of microRNAs enriched in LSC by RT-qPCR (reverse transcription followed by quantitative PCR) in additional CML patients; bioinformatic analysis of their associated targets revealed an enrichment of multiple metabolic pathways, suggesting that microRNAs may be important mediators of LSC dysregulated metabolism.

## Results

### Global patterns in the miRNome of LSC and CML HSC

We isolated highly enriched LSC and HSC fractions from CML patients at diagnosis or from HD, based on a combination of cell surface markers (CD34, CD38, CD45, CD26) and flow cytometry parameters (FSC, SSC) (see Supplementary Fig. S1). Some patients showed no clear separation of CD26^-^ and CD26^+^ populations (see Supplementary Fig. S2): in those cases, both fractions included leukemic *BCR-ABL1*^+^ cells, therefore, we only used CD26^+^ fraction. Purity was assessed by *BCR-ABL1* mRNA detection in CFU-derived colonies (see Supplementary Fig. S2). We extracted total RNA containing the small RNA fraction (<200nt) from sorted cells; given that individual patient-derived fractions had low yields of RNA, samples from different patients or HD were pooled before preparation of libraries for small RNA-NGS(CML LSC CD34^+^CD38^-^CD26^+^, CML HSC CD34^+^CD38^-^CD26^-^, HD HSC CD34^+^CD38^-/dim^, HD progenitors CD34^+^CD38^+^). More than 1,000 (≥1 count) or 600 (≥10 counts) different microRNAs were detected in each fraction, with high abundance of a few specific microRNAs: top-10 most abundant microRNAs represented 52-65% of total microRNAs in CML and HD (Fig. 1a). Most (>80%) microRNAs dysregulated (GFOLD≥|1|) in LSC and HSC from CML patients had decreased levels compared to primitive (CD34^+^CD38^-/dim^) cells from HD, suggesting a global pattern of microRNA downregulation (Fig. 1b).

**Fig. 1.**
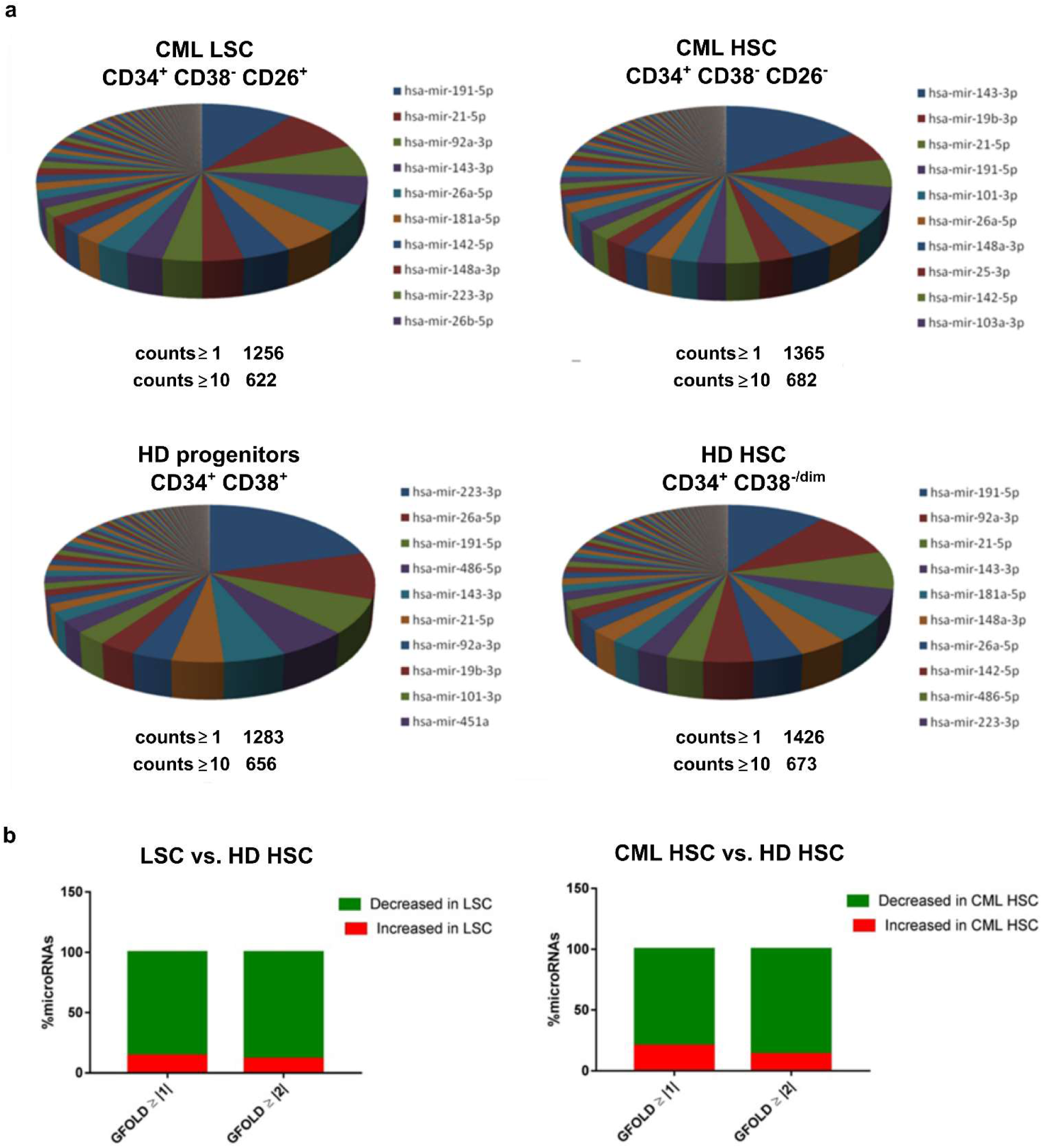
Global patterns in the miRNome of LSC and CML HSC. a: Pie chart representing the relative abundance of each microRNA in each fraction assessed by small RNA-NGS. The total number of different microRNAs (with at least 1 or 10 counts) is detailed below the pie chart. The names of the top-10 most abundant microRNAs in each fraction are detailed. b: Global decrease in microRNA levels in CML primitive cells compared to HD HSC. Most microRNAs dysregulated in both LSC and HSC from CML samples had decreased levels (GFOLD ≥ |1| or GFOLD ≥ |2|) compared to HD HSC.

### microRNAs enriched in LSC vs. HSC from CML patients and HD

Differential expression of microRNAs was assessed by calculation of a GFOLD value, which is a robust fold-change parameter that considers both the absolute number and the relative difference in microRNA levels between samples. With a cut-off value of GFOLD ≥ |1|, we found 120 microRNAs dysregulated between LSC and putative HSC from CML patients; and 46 microRNAs between CML LSC and HSC of HD. The intersection of both lists resulted in 16 microRNAs (see Supplementary Table S2 and Fig. 2a). *In silico* analysis of both predicted and experimentally validated targets resulted in clusters of functionally related microRNAs that correlated with increased or decreased levels in LSC vs. putative HSC from CML patients, suggesting the existence of mechanisms that regulate microRNAs levels in a coordinated fashion (Fig. 2c). We observed enrichment in pathways related to fatty acid metabolism/biosynthesis, Hippo signaling, *adherens* junction, proteoglycans in cancer, and extracellular matrix-receptor interaction (Fig. 2c). In order to evaluate possible bias in the results due to the inclusion of experimentally validated microRNA-mRNA interactions, we performed an identical pathway enrichment analysis using two lists of 16 randomly selected microRNAs, among those that were detectable, but that did not vary (GFOLD=0) among LSC and CML HSC fractions. This analysis resulted in fewer pathways, microRNAs and genes in both lists of microRNAs (see Supplementary Table S3), supporting the validity of results obtained for dysregulated microRNAs. Surprisingly, most (7 out of 8) microRNAs with decreased levels in LSC belong to a genomic cluster located in region 14q.32 (*DKL1/DIO3* locus). Moreover, inspection of microRNAs and snoRNAs from this locus revealed that 18 additional microRNAs and 5 snoRNAs had decreased levels in LSC vs. HSC from HD (Fig. 3 and Supplementary Table S4). Given that this region contains imprinted genes^23^, this result suggests the presence of a mechanism of epigenetic silencing in LSC.

**Fig. 2.**
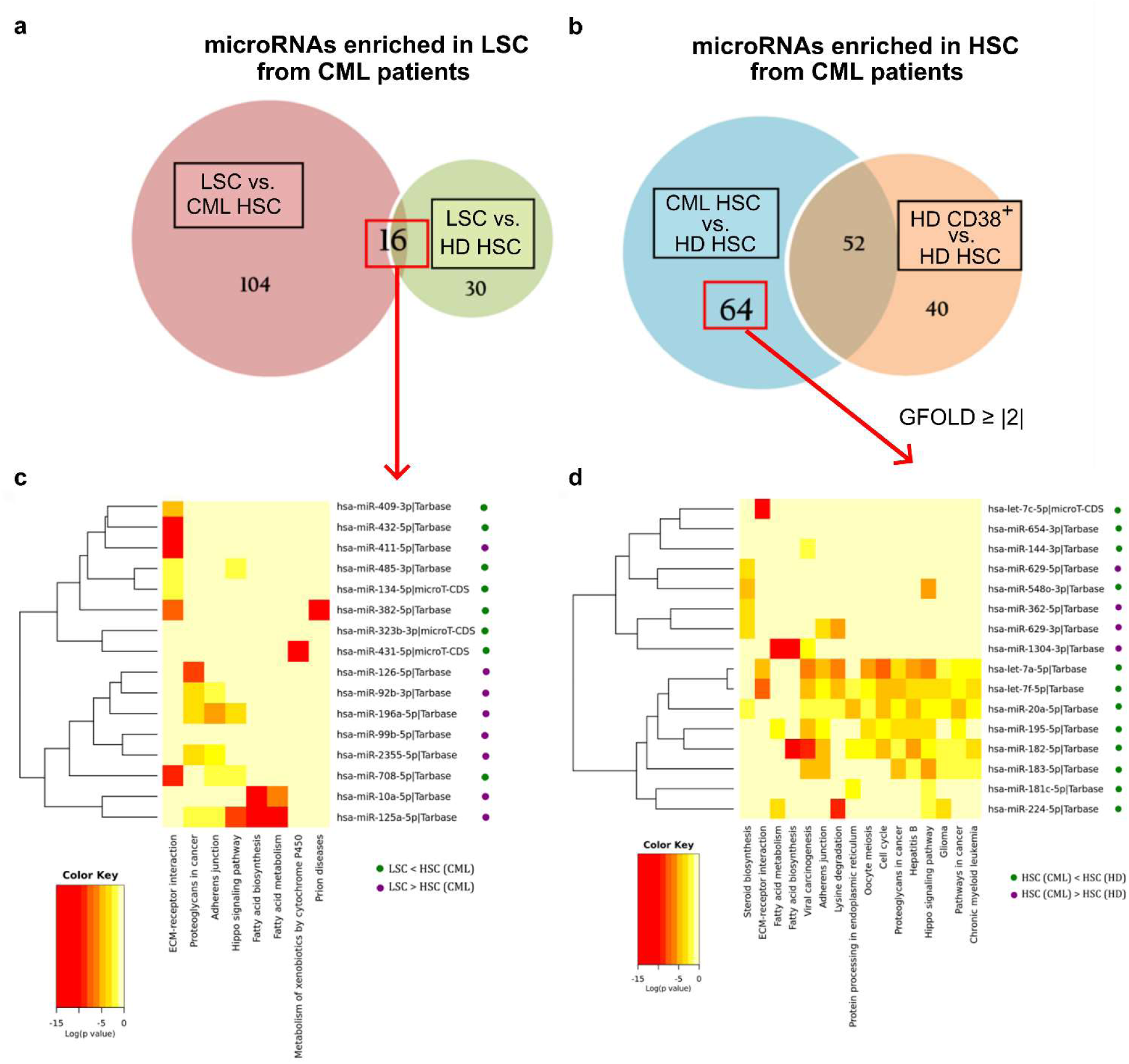
Differential expression and pathway analysis of small-RNA NGS data. a, b: Number of microRNAs with GFOLD≥|1| in each comparison. Selected microRNAs are indicated in red. c, d: Heatmaps of KEGG molecular pathways associated with predicted and experimentally validated targets of microRNAs dysregulated in LSC (c, miRPath, pathway union) or HSC from CML patients (d, miRPath, pathway union). Green and purple dots refer to the relative abundance of each microRNA in each population.

**Fig. 3.**
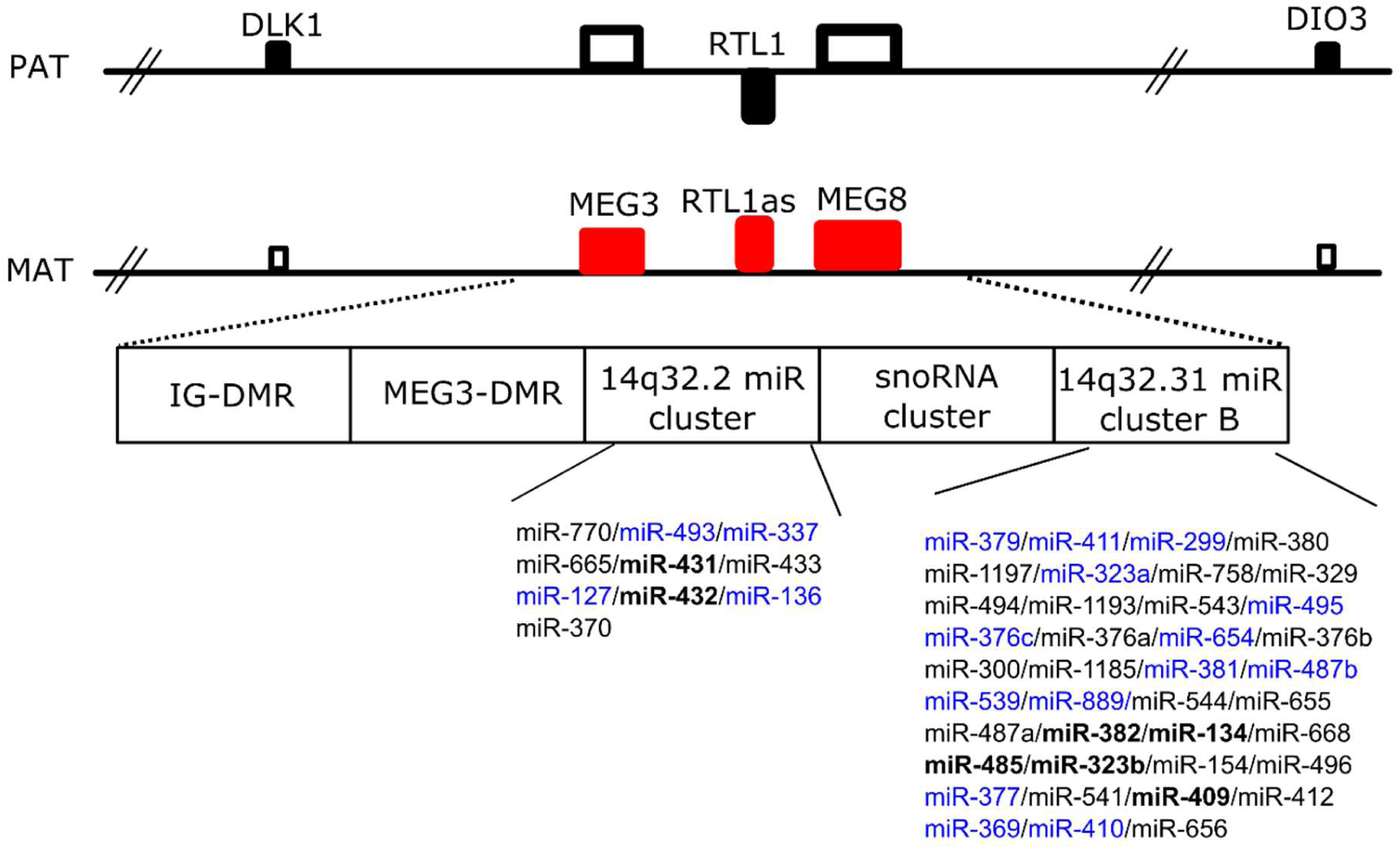
Schematic representation of the genomic 14q.32 region. This region includes coding genes with paternal (PAT) imprinting (*DLK1, RTL1, DIO3*), non-coding genes with maternal (MAT) imprinting (*MEG3*, anti*-RTL1, MEG8*), two clusters of microRNAs, and one cluster of snoRNAs. microRNAs in bold showed decreased levels in LSC vs. both HD HSC and CML HSC; while microRNAs in blue showed decreased levels in LSC vs. HD HSC but did not vary compared to CML HSC. IG: intergenic. DMR: differentially methylated regions.

### microRNAs in HSC from CML patients show a dysregulated miRNome despite the absence of BCR-ABL1

Based on the hypothesis that HSC present in CML patients are not equivalent to HSC in HD, we compared microRNAs between both fractions. We found 64 microRNAs significantly dysregulated (Fig. 2b); further selection (GFOLD ≥ |2|) resulted in a list of 16 microRNAs (see Supplementary Table S5). It is important to clarify that HSC from HD were sorted using a less strict gating of CD38-negative cells (resulting in a CD38^-/dim^ population), because we obtained very low yields from individual samples. Therefore, in order to exclude differentially expressed microRNAs related to the inclusion of a CD38^dim^ population in HD, we excluded microRNAs that were differentially expressed between CD38^-/dim^ and CD38^+^ fractions from HD, under the assumption that some of these microRNAs would be related to the process of hematopoietic differentiation (Fig. 2b). *In silico* analysis of both predicted and experimentally validated targets resulted in a cluster of microRNAs with decreased levels in HSC from CML patients, which were enriched in proliferative pathways such as viral carcinogenesis, cell cycle, hepatitis B, pathways in cancer, and chronic myeloid leukemia (Fig. 2d), among other pathways that were also present in LSC (Hippo signaling, proteoglycans in cancer, fatty acid metabolism, extracellular matrix-receptor interaction). These results suggest that putative HSC from CML patients are either altered by extrinsic factors (i.e. a niche altered by coexistence with leukemic cells, or even the transfer of microRNAs from neighbor cells), and/or that they are pre-leukemic neoplastic stem cells and thus harbor early, BCR-ABL-independent genetic or epigenetic alterations that affect microRNA levels (i.e. mutations in microRNA-processing machinery).

### Validation by RT-qPCR in a new cohort of CML patients and HD

As other techniques, NGS is not free of intrinsic bias, mainly related to library preparation, platform used for sequencing, and data analysis^24^. In addition to perform a technical validation, we aimed to validate NGS results in a new cohort of patients and HD using RT-qPCR. We performed a multiplex RT step that allowed us to measure individual microRNA levels using very low inputs of RNA; therefore, pooling of samples from different patients or HD was not necessary, and we could assess intra-group variability. We evaluated the following fractions from CML patients or HD samples: CML LSC (CD34^+^CD38^-^CD26^+^), CML HSC (CD34^+^CD38^-^CD26^-^), CML progenitors (CD34^+^CD38^+^), HD HSC (CD34^+^CD38^-/dim^), HD progenitors (CD34^+^CD38^+^).

We selected six microRNAs upregulated in LSC vs. HSC from CML patients (miR-125a-5p, miR-10a-5p, miR-126-5p, miR-92b-3p, miR-196a-5p, miR-2355-5p), and measured their levels in LSC, HSC and progenitor fractions from six CML patients, and also in HSC and progenitor fractions from four HD. We also included four additional small RNAs: one potential “novel” microRNA that emerged from NGS-data (“novel-3”), and three additional microRNAs which were of interest in this population according to previous references (miR-let-7a-5p, miR-132-3p, miR-182-5p). Purity of fractions was assessed by RT-qPCR of *BCR-ABL1* in RNA isolated from sorted cells (see Supplementary Fig. S3). miR-2355-5p was not detected in most fractions, therefore we excluded it from posterior analysis. We detected significant differences among fractions for miR-125a-5p, miR-10a-5p, miR-126-5p, miR-92b-3p, and miR-196a-5p (global p-value < 0.05; linear mixed-effects model) (Fig. 4). We could not detect global differences in the levels of “novel-3”, miR-let-7a-5p, miR-132-3p, and miR-182-5p (global p-value > 0.05; linear mixed-effects model) (see Supplementary Fig. S4). The trend of change was maintained between NGS and RT-qPCR for miR-125a-5p, miR-196a-5p, and miR-92b-3p (increased levels in LSC vs. HSC from CML patients) (see Supplementary Fig. S4). miR-126-5p and miR-10a-5p displayed opposite trends in NGS and RT-qPCR (see Supplementary Fig. S5). CML progenitors were only measured by RT-qPCR; interestingly, miR-10a-5p and miR-125a-5p were decreased in CML progenitors vs. CML primitive cells, while miR-92b-3p levels were increased (see Supplementary Fig. S6).

**Fig. 4.**
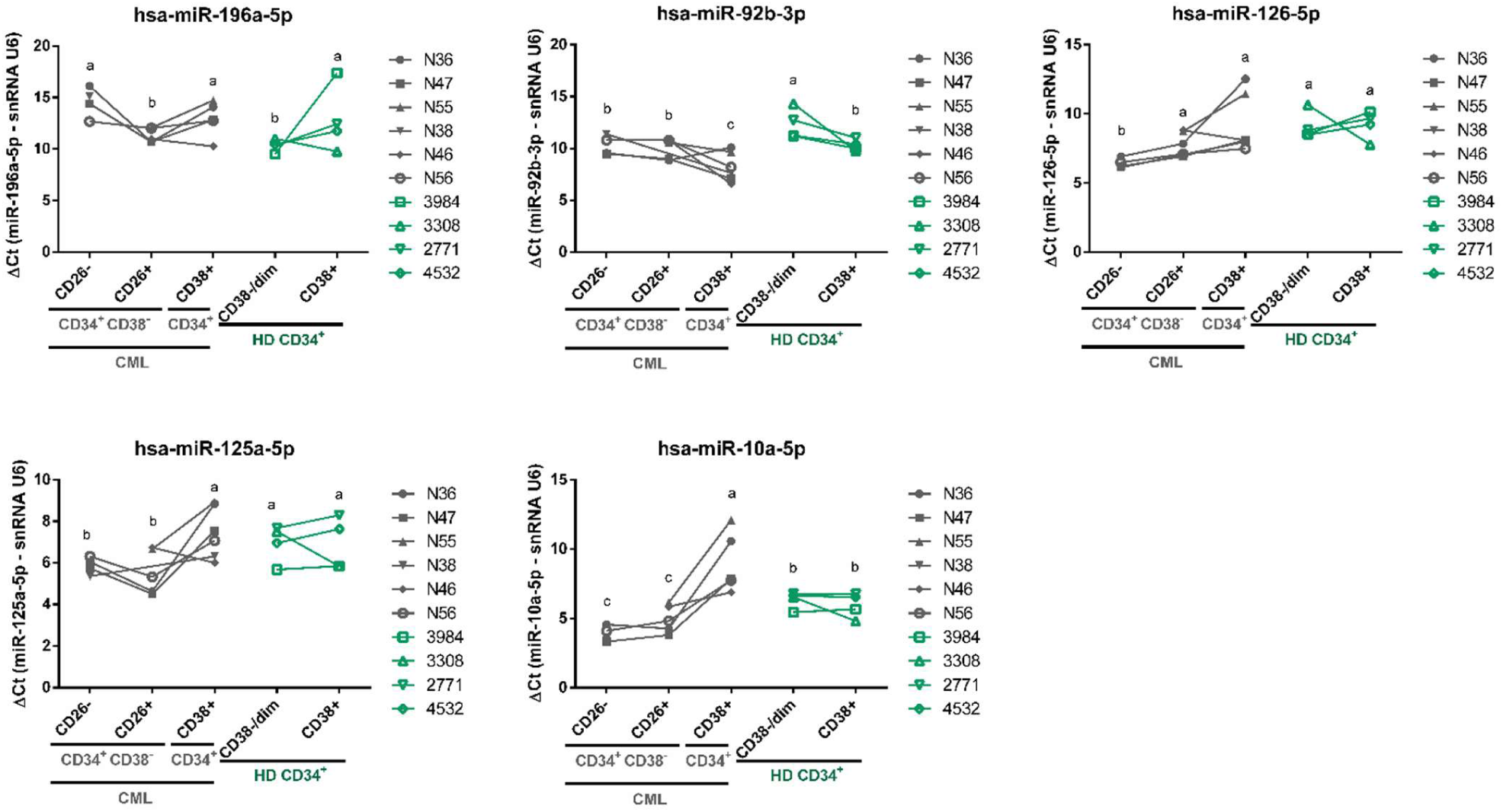
Validation of microRNAs by RT-qPCR in a new cohort of CML and HD samples. Results are expressed as ΔCt = Ct (microRNA) - Ct (snRNA U6). Each dot is the mean of technical duplicates from each patient or HD. CML samples are represented in grey symbols, and HD samples in green symbols. Lines connect different fractions from the same patients or HD. Different letters indicate statistically significant differences (linear mixed-effects model, *a posteriori* comparison, global α = 0.05)

*In silico* analysis of both predicted and experimentally validated targets of miR-125a-5p, miR-10a-5p, miR-92b-3p and miR-196a-5p by different bioinformatics tools (see Methods section) resulted in enrichment in KEGG (Kyoto Encyclopedia of Genes and Genomes) pathways (5 out of 10 in ChemiRs; 3 out of 7 in miRPath) and Gene Ontology (GO) terms (4 out of top-10 in ChemiRs; 3 out of top-10 in miRPath) related to metabolic processes, including lipid, sugar, and nitrogen compound metabolism (Table 2). We searched for possible targets included in hallmark gene sets^25^ of fatty acid metabolism, oxidative phosphorylation and glycolysis. The search of potential targets included predicted and experimentally validated microRNA-mRNA interactions, and the intersection with published lists of mRNAs^16^ and proteins^18^ dysregulated in CML precursor/primitive fractions (Fig. 5). This analysis reduced the number of potential targets from 1,659 to 16 genes, some of them with experimental evidence of altered levels in CML LSC or CD34^+^ cells.

**Table 1.** CML and HD samples used for small RNA-NGS and validation by RT-qPCR. The number of events refers to the number of sorted cells obtained from each fraction. NE*: fractions not evaluated (pattern 3). ND: not done

| Code | Type | Sex | Age | HSC events | LSC events | Progenitors events |
| --- | --- | --- | --- | --- | --- | --- |
| CML samples used for small RNA-NGS (pooled) |  |  |  |  |  |  |
| N26 | BM | M | 24 | 6896 | 1857 | ND |
| N33 | PB | M | 54 | 1859 | 0 | ND |
| IM | BM | M | 22 | NE* | 3046 | ND |
| HD samples used for small RNA-NGS (pooled) |  |  |  |  |  |  |
| 2891 | buffy coat | M | 36 | 1578 |  | 7624 |
| 2890 | buffy coat | F | 31 | 1316 |  | 13724 |
| 2810 | PB | F | 50 | 156 |  | ND |
| 3060 | PB | F | 54 | 181 |  | ND |
| 3308 | buffy coat | F | 37 | 1700 |  | 13000 |
| CML samples used for validation by RT-qPCR |  |  |  |  |  |  |
| N36 | PB | M | 56 | 2857 | 2200 | 37272 |
| N47 | PB | M | 58 | 1128 | 4504 | 96333 |
| N55 | PB | M | 25 | NE* | 9276 | 19861 |
| N38 | PB | M | 31 | 592 | 0 | 2300 |
| N46 | PB | F | 38 | NE* | 3360 | 61428 |
| N56 | PB | M | 55 | 4267 | 2910 | 291653 |
| HD samples used for validation by RT-qPCR |  |  |  |  |  |  |
| 3984 | buffy coat | F | 36 | 1117 |  | 1779 |
| 3308 | buffy coat | F | 37 | 1682 |  | 2027 |
| 2771 | buffy coat | M | 53 | 4460 |  | 9539 |
| 4532 | buffy coat | F | 57 | 2314 |  | 13256 |

**Table 2.** Top-ten results for KEGG molecular pathways and GO terms analysis, associated to hsa-miR-10a-5p, hsa-miR-125a-5p, hsa-miR-92b-3p or hsa-miR-196a-5p. Processes and terms related to cell metabolism are highlighted in bold.

| KEGG pathways (ChemiR) | KEGG pathways (miRPath-Tarbase) | GO terms (biological process) (ChemiR) | GO categories (biological process) (miRPath-Tarbase) |
| --- | --- | --- | --- |
| 1. **Metabolism** | 1. **Fatty acid biosynthesis** | 1. **Canonical glycolysis** | 1. **Nucleobase-containing compound metabolic process** |
| 2. Bladder cancer | 2. **Fatty acid metabolism** | 2. **Cellular glucose homeostasis** | 2. Membrane organization |
| 3. Elongation arrest and recovery | 3. Hippo signaling pathway. | 3. Negative regulation of epithelial to mesenchymal transition | 3. Mitotic cell cycle |
| 4. **Fructose and mannose metabolism** | 4. **Lysine degradation.** | 4. Regulation of transcription involved in G2/M transition of mitotic cell cycle | 4. **mRNA metabolic process** |
| 5. **Biosynthesis of unsaturated fatty acids** | 5. *Adherens* junction | 5. Skeletal system morphogenesis | 5. Cellular component assembly |
| 6. **Galactose metabolism** | 6. Proteoglycans in cancer | 6. **ncRNA metabolic process** | 6. Fc-epsilon receptor signaling pathway |
| 7. Circadian rhythm - mammal | 7. Chronic myeloid leukemia | 7. Release of cytochrome c from mitochondria | 7. **Cellular protein metabolic process** |
| 8. **Alpha-linolenic acid metabolism** |  | 8. RNA splicing, via transesterification reactions | 8. Neurotrophin TRK receptor signaling pathway |
| 9. Translocation of SLC2A4(Glut4) to the plasma membrane |  | 9. **Carbohydrate phosphorylation** | 9. Cellular protein modification process |
| 10. Protein export |  | 10. Chaperone-mediated protein folding | 10. Response to stress |

**Fig. 5.**
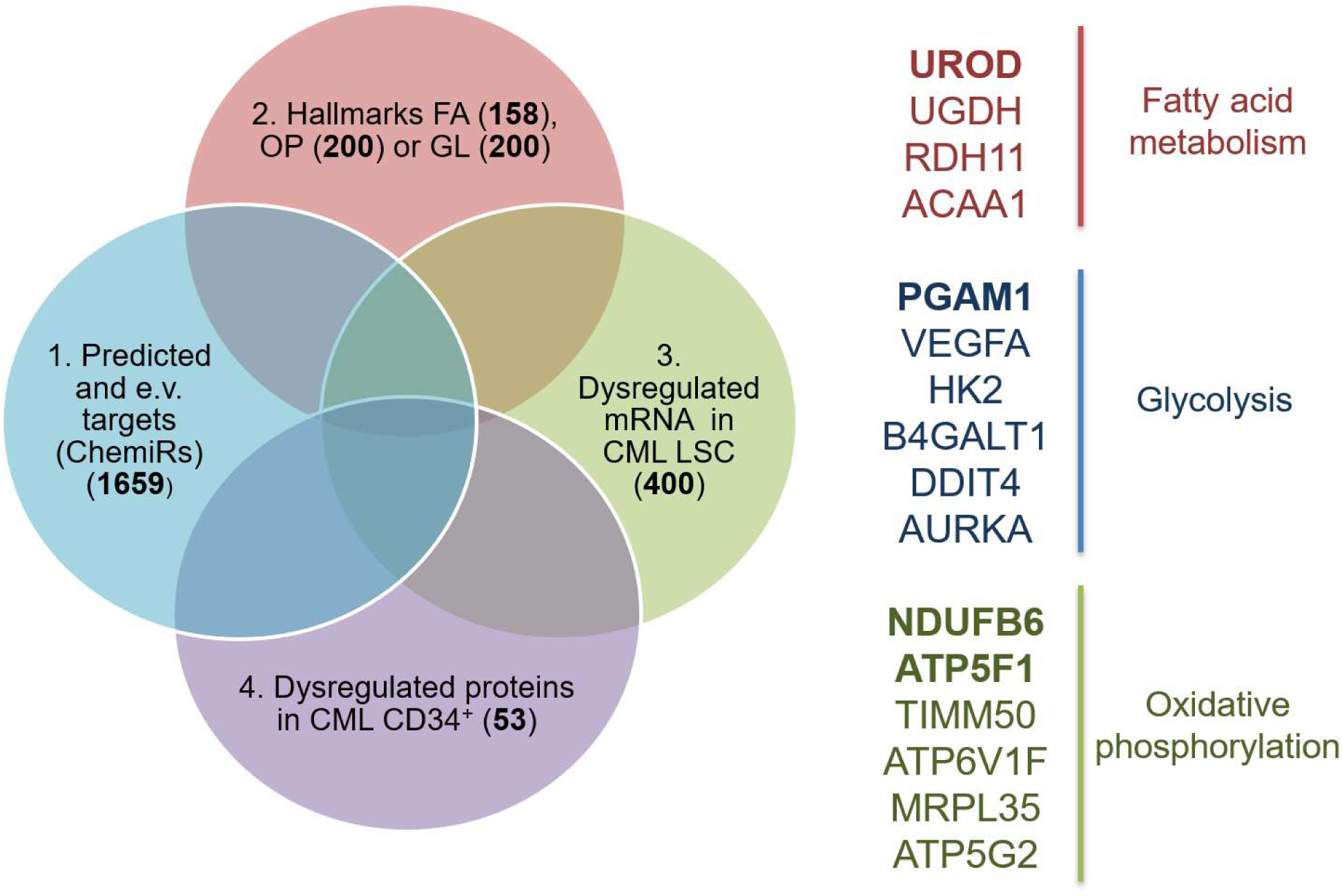
Selection of potential targets of validated microRNAs. The search of potential targets included predicted and experimentally validated (e.v.) microRNA-mRNA interactions for miR-10a-5p, miR-92b-3p, miR-196a-5p, and miR-125a-5p (list 1), intersected with hallmark gene sets “FA: fatty acid metabolism”, “OP: oxidative phosphorylation”, or “GL: glycolysis” (list 2), and with published lists of mRNAs^16^ (list 3) or proteins^18^ (list 4) dysregulated in CML precursor/primitive fractions. This analysis reduced the number of potential targets from 1,659 to 16 genes, some of them with experimental evidence of altered levels in CML LSC and/or CD34^+^ cells (genes in bold). The number of genes in each list is indicated in bold.

## Discussion

To our knowledge, this is the first description of the miRNome of CML LSC. In these analyses, major differences were found when analyzing *BCR-ABL1*^+^ and *BCR-ABL1*^-^ primitive cells in newly diagnosed patients. In addition, we detected several major differences in the miRNome pattern when comparing CML LSC with HSC of HD. First, we observed a global downregulation of microRNAs in primitive CML cells in comparison with HSC of HD. Second, differentially expressed microRNAs showed covariation according to their potential targets, suggesting that mature levels of functionally related microRNAs are (dys)regulated by common mechanisms. Third, compared to HSC from HD, we detected decreased levels in LSC of microRNAs and snoRNAs belonging to a genomic cluster located in chromosome 14 (14q.32). Fourth, a high number of microRNAs were differentially expressed between putative HSC from CML patients and HSC from HD, suggesting an altered phenotype of the “normal” HSC fraction in CML patients. Finally, we confirmed by RT-qPCR that the levels of miR-125a-5p, miR-10a-5p, miR-92b-3p and miR-196a-5p were altered in CML primitive and progenitor fractions and were enriched in metabolism-related targets.

Global downregulation of microRNAs in cancer has been reported in different tumors^26^. Multiple mechanisms have been described to explain microRNA dysregulation in cancer, including genomic structural variations (i.e. deletions, amplification or translocations), alterations in oncogenes or tumor suppressor genes that regulate microRNA transcription, and epigenetic changes (such as hypermethylation of microRNA promoters). Defective microRNA processing machinery has been described as well: for example, altered expression and function of the Microprocessor components (i.e. Drosha, DGCR8), and dysregulation of the complex that mediates pre-microRNA export from the nucleus^27^. Recent work by Mori *et al* described a link between dysregulation of Hippo signaling pathway in cancer, and mature microRNA depletion^28^. Interestingly, global microRNA loss was shown to enhance tumorigenesis^29^. In CML, global microRNA depletion in patient samples has not been reported so far. In the work of Zhang *et al*, they showed, in K562 cells and CML CD34^+^ cells, that BCR-ABL1 can affect the export of miR-126 precursors from the nucleus to the cytoplasm, through phosphorylation of SPRED1, a negative regulator of RAS superfamily proteins, interfering with Ran-exportin 5-RCC1 complex^30^. Interestingly, this effect was reversible by treatment with Nilotinib. However, in our work, we observed decreased levels of mature microRNAs in both *BCR-ABL1*^*+*^ and *BCR-ABL1*^*-*^ primitive (CD34^+^CD38^-^) cells compared to HSC from HD, suggesting a BCR-ABL1-independent mechanism.

Clustering of microRNAs dysregulated in LSC according to their target-related pathways correlated with their increased or decreased levels in LSC vs. their *BCR-ABL1*^-^ counterparts (CML HSC), suggesting the existence of mechanisms of coordinated regulation. MicroRNAs can belong to families in which members are evolutionary related, therefore they share regions of common sequences, and can regulate similar or related targets. In this context, miR-125a and miR-10a belong to the miR-10/miR-100 family, and we found a significant positive correlation of both microRNAs in samples evaluated by RT-qPCR (r (Pearson)=0.85; p=5.7×10^−7^). This suggests that future studies aimed at evaluating the functional relevance of microRNAs dysregulated in this system should take into consideration possible functional redundancy between related microRNAs. In fact, knockout experiments of microRNAs belonging to the same family have shown partially redundant effects on mice^31^. Therefore, the combination of individual and simultaneous knockdown of correlated microRNAs would be an ideal approach.

In humans, the *DKL1/DIO3* locus, at the 14q.32 region, contains the paternally expressed genes *Delta-like 1 homolog* (*DLK1), Retrotransposon-like 1* (*RTL1)*, and *Iodothyronine deiodinase 3* (*DIO3)*, and the maternally expressed genes *MEG3, MEG8*, and anti-sense *RTL1. DLK1* is a non-canonical Notch ligand, and seems to be involved in developmental processes, such as branching morphogenesis and terminal differentiation^32^. *RTL1* is a key gene in placental development and evolution^33^, whereas *DIO3* protects developing tissues from excessive amounts of thyroid hormone^34^. *MEG3* and *MEG8* are long intergenic RNAs; *MEG3* has been found deregulated in several types of tumors, and it is believed to function as a tumor-suppressor gene through interactions with p53^35^, and to participate in epigenetic regulation by interacting with chromatin modifying complexes such as PRC2^36^. The largest mammalian cluster of microRNAs, together with a cluster of snoRNAs, are included in the maternally expressed strand. We observed a downregulation of microRNAs and snoRNAs from the 14q.32 cluster in the NGS cohort, suggesting a process of epigenetic silencing of the corresponding allele in LSC. Interestingly, chromosome 14q acquired uniparental disomy (aUPD) is one of the most common abnormalities associated with clonal hematopoiesis in elderly individuals; a recent report identified that *MEG3-DLK1* locus at 14q.32 is the primary target of aUPD^37^, suggesting that this phenomenon would not be unique to CML pathogenesis. On the other hand, *MEG3* was shown to be downregulated in CML chronic phase samples. Furthermore, patients in advanced phase and blast crisis showed further decreased levels of *MEG3*^38^. An exciting discovery performed in a murine model was the observation that non-coding RNAs (including microRNAs) from this locus maintain fetal liver and adult long-term repopulating HSCs (LT-HSCs) through the suppression of the PI3K-mTOR pathway, which results in inhibition of mitochondrial biogenesis and metabolic activity^39^. In the light of the recently described increase in oxidative metabolism in CML LSC^19^, is there an association between the loss of expression of microRNAs from DKL1/DIO3 locus in LSC and their altered metabolome? Furthermore, could be the increase in mitochondrial activity in CML LSC related to their loss of quiescence^17^?

The role of cell-extrinsic factors in the development of hematological malignancies is an exciting field. A recent study on LSC and HSC from CML patients revealed that the transcriptional profile of HSC, assessed by single-cell analysis, was more informative than LSC to allow clustering of cells derived from TKI-non-responder and responder patients^16^. In our study, a great number of microRNAs were dysregulated between HSC from CML patients and HSC from HD. The fact that HSC from CML patients showed an altered miRNome could be attributed to cell-autonomous (i.e. genetic or epigenetic alterations) and/or extrinsic factors, such as an altered microenvironment due to the presence of high numbers of leukemic cells, where microRNAs trafficking among different cell types may act as signals coming from CML cells (i.e. through extracellular vesicles)^40^. Therefore, it would be of interest to assess whether the miRNome of CML HSC is restored upon TKI treatment in all patients. An alternative explanation would be that some or even most of the putative HSC in our CML patients were indeed pre-leukemic neoplastic stem cells (pre-L-NSC)^10,13^. In CML such early pre-L-NSC may express BCR-ABL1-independent molecular and epigenetic lesions and may be negative for CD26. Finally, it cannot be excluded that some of the genetic/molecular material was transferred from CML cells to normal cells or pre-L-NSC and thereby introduced oncogenic signaling pathways leading to an altered miRNome.

miR-126 has been extensively studied because of its functional relevance in the vascular system. Regarding its genomic location and transcriptional regulation, it is intragenic (intronic), and its host gene (EGFL7) is a peptide present in endothelial cells. In the hematopoietic system, miR-126 has been described as a regulator of stem-cell properties, and it is present at high levels in the most primitive hematopoietic cell fraction (CD34^+^CD38^-^CD45RA^-^CD49f^+^)^41^. In CML, mature miR-126-3p levels are reduced in LSC in a BCR-ABL1-kinase activity-dependent manner. However, there are no reports of miR-126-5p in this system^30^. Interestingly, miR-126-3p levels in our NGS-cohort were lower than those of mi-126-5p, and therefore it was not selected for further validation. However, they were lower in LSC compared to CML HSC and HD HSC (0.59 and 0.35-fold-change, respectively), in agreement with the data reported by Zhang et al^30^. Additionally, the authors showed that miR-126-3p can be transferred from BM-derived endothelial cells to primitive hematopoietic cells by extracellular vesicles. Therefore, the results obtained in this work suggest the prospective analysis of miR-126-5p and miR-126-3p of paired samples of BM and PB-derived LSC and HSC.

Bioinformatic tools for microRNA analysis are part of a growing field. We first evaluated possible functional relevance of microRNAs of interest by analyzing common molecular pathways associated with their possible targets. However, given the pleiotropic effects of microRNAs, which can regulate several targets simultaneously, target prediction presents high rates of false positives. Therefore, we filtered target predictions by considering only those microRNA-mRNA interactions that had been experimentally validated, which included non-canonical interactions that are usually excluded from target prediction algorithms. This strategy resulted in the enrichment of targets related to metabolic processes. In the light of recent descriptions of an altered metabolome in LSC from CML^16^, our results highlight the potential of microRNAs to reveal biological patterns. Previous reports have shown that microRNA profiles are more informative than mRNA profiles to classify human cancers^42^, possibly because the network of microRNAs has a lower dimensionality than the network of mRNAs. Results obtained from our *in silico* analysis serve as a guide for future functional studies that evaluate the link between microRNAs and metabolic dysregulation in LSC. Given that HSC require a delicate regulation of cell metabolism in order to maintain their phenotype^43^, these results open exciting questions about the role of microRNAs in the modulation of CML LSC. Finally, LSC in CML patients at diagnosis comprise a heterogeneous fraction according to surface markers and mRNA levels^12,16^; however, there are no reports on the involvement of microRNAs in the definition of these populations. In this context, does the pattern of microRNAs vary among individual LSC and HSC? Given that microRNAs have been proposed as regulatory molecules able to confer robustness to a biological process, by fine-tuning of mRNA and protein levels under a specific context, it would be exciting to explore patterns of microRNAs at the single-cell level in LSC and HSC from CML patients, in order to further understand the biological properties of these heterogeneous fractions.

## Methods

### 1. Patient samples

The project was approved by the Institutional Review Board, at Instituto Alexander Fleming (Buenos Aires, Argentina). All procedures involving human participants were in accordance with the ethical standards of the institutional research committee and with the 1964 Helsinki declaration and it later amendments. All patients and healthy donors gave written informed consent. Bone marrow (BM) or peripheral blood (PB) samples were obtained from newly diagnosed, untreated CML patients in chronic phase. Patient samples used for library preparation for small-RNA-NGS and validation by RT-qPCR are listed in Table 1. Mononuclear cells (MNC) were isolated by density-gradient centrifugation (Ficoll-Paque PLUS, GE Healthcare Life Sciences) for 30 minutes at 400 x*g*, followed by one wash in phosphate buffered saline (PBS, GIBCO), a red cells lysis step, and a low-speed centrifugation step (12-15 minutes at 200 x*g*) for removal of the platelet-rich fraction. Up to 2×10^8^ MNC were used for isolation of CD34^+^ cells.

### 2. Isolation of CD34^+^ cells

In order to enrich for stem and progenitor cells, we performed a positive selection using CD34 MicroBeads (Miltenyi Biotech), according to manufacturer’s instructions. The CD34^+^ fraction was immediately used or cryopreserved in 1 mL of freezing medium (see Supplementary methods).

### 3. CFU assay for assessment of purity in sorted fractions

Between 250-500 CD34^+^ cells were directly sorted into 250 µL of enriched methylcellulose (Methocult H4435 Medium, Stem Cell Technologies, Vancouver, Canada), and then plated into p35 culture dishes containing a final volume of 1.1 mL of enriched methylcellulose. Cells were incubated at 37°C in a humid chamber. After 14-18 days, pools of 4-6 colonies (CFU-GM, BFU-E and mixed CFU-GEMM) were plucked from methylcellulose, resuspended in 500 µL of Roswell Park Memorial Institute - 1640 medium (RPMI-1640, GIBCO), and centrifuged. Cells were resuspended in 100 µL of lysis solution (RNAqueous-Micro Kit, Ambion), and kept at -20°C until RNA extraction was performed. Total RNA was extracted following manufacturer’s instructions, and *BCR-ABL1* mRNA was measured by RT-qPCR (see Supplementary methods).

### 4. Isolation of LSC and HSC by FACS

Total number of cells used for FACS varied according to the yield of each sample. CD34^+^ cells or MNC from CML patients or HD were incubated with the following antibodies: 5 µL CD45-PerCP (2D1, BD Biosciences), 2.5 µL CD34-FITC (AC136, Miltenyi Biotech), 2.5µL CD38-PE (IB6, Miltenyi Biotech), and 15 µL CD26-APC (FR10-11G9, Miltenyi Biotech), in a final volume of 100 µL of MACS buffer, for 15 minutes at room temperature. Cells were washed once with 1 mL of PBS (GIBCO) and resuspended in 300 µL of PBS. Sorting was performed in a FACS Aria II cytometer (BD Biosciences), located at Facultad de Ciencias Exactas y Naturales, Universidad de Buenos Aires (Buenos Aires, Argentina). Setting of positive and negative gates for CD38, CD34, and CD26 was performed on the CD45^low^ population; therefore, isotype control tube included 2.5 µL Mouse IgG2a-FITC (Miltenyi Biotech), 2.5 µL Mouse IgG2a-PE (Miltenyi Biotech), 15 µL Mouse IgG2a-FITC (Miltenyi Biotech), and 5 µL CD45-PerCP. In order to avoid electronic aborts that could affect the purity of sorted fractions, the parameter “window extension” was set to zero. Other parameters included 70µm nozzle, and “purity”. Cells were collected in aseptic conditions, directly into 100 µL of lysis buffer for RNA extraction (RNAqueous-Micro Kit, Ambion), in RNAse-free 200 µL tubes, or in enriched methylcellulose for assessment of purity. Flow-cytometry data were analysis was performed with BD FACSDiva (version 6.1.3) and FlowJo (version 7.6.2) software.

Total RNA containing small RNAs (<200nt) was extracted following the protocol from RNAaqueous-micro kit (Ambion) with a slight modification: 125 µL of EtOH 100% were added to the lysate and vortexed; the rest of the protocol was performed according to manufacturer’s instructions. RNA elution was performed twice (9 µL each) using pre-warmed distilled water (75°C). RNA was kept at -80°C. Quality and conservation of the small RNA fraction were assessed with Agilent 2100 Bioanalyzer (total RNA Nano kit), at Fundación Instituto Leloir (Buenos Aires, Argentina).

### 5. Concentration of pooled samples for small RNA-NGS

RNAs extracted from different samples were combined in order to increase RNA yield before NGS-library preparation. After mixing, RNA was freezed at -80°C, and transported from Argentina to Brazil in dry ice. Samples were concentrated using a vacuum centrifuge and resuspended in 7 µL of distilled water (5 minutes at 50°C). Quantification of RNA was performed using Qubit 2.0. The entire content was used for library preparation (<140 ng for LSC, CML HSC and HD HSC fractions, and 210 ng for HD progenitor fraction).

### 6. Preparation of libraries for small RNA-NGS in HiSeq 2500 (Illumina)

Libraries from each pool of samples (CML LSC CD34^+^CD38^-^CD26^+^, CML HSC CD34^+^CD38^-^CD26^-^, HD HSC CD34^+^CD38^-^ /dim, HD progenitors CD34^+^CD38^+^) were prepared using Truseq Small RNA kit (Illumina), following manufacturer’s instructions (15 PCR cycles). The protocol is based on the selective ligation of RNAs with free 3’OH and 5’phosphate ends, resulting from precursor cleavage during small RNA biogenesis^44^. Therefore, other small RNAs besides microRNAs are included in the library: fragments of tRNAs, small nucleolar RNAs (snoRNAs), small nuclear RNAs (snRNAs), and piwiRNAs. Estimated size of the libraries was 147-157bp, which were purified by band excision after polyacrylamide denaturing gel electrophoresis (Novex 6% TBE, Invitrogen). Quantification of libraries was performed by qPCR (KAPA SYBR, Roche Life Sciences). Libraries were concentrated before sequencing by vacuum centrifugation. Single-end sequencing was performed on HiSeq 2500 (Illumina) at Instituto Nacional de Câncer (Rio de Janeiro, Brazil).

### 7. Bioinformatic analysis of small RNA-NGS

Low quality lectures were filtered (fastq_quality_filter; >80% reads with Q>20), and contaminant sequences were removed (3’ and 5’ adaptors, indexes). Identification of known microRNAs was performed with Chimira^45^; raw microRNA counts from each pool of samples is available (see Supplementary Data 2.xlsx). Differential expression analysis was performed using GFOLD algorithm (c=0.01), which is especially suited for experiments without biological replicates, after mapping against a database of snoRNA/miRNA (HISAT2)^46^. Complete results from GFOLD analysis are available (see Supplementary Data 1.xlsx). Analysis of potential targets and related pathways was performed using miRPath (*Diana tools*)^47^, and ChemiRs^48^. The parameters used for miRPath analysis were: “*KEGG analysis*”, *Tarbase* (database of experimentally validated interactions), or *microT-CDS* in those cases with no experimental evidences, “*Pathway union*”, “*p-value threshold:* 0.001”, “*MicroT threshold*: 0.8”, “*Enrichment analysis method= Fisher’s exact test (hypergeometric distribution)*, “*FDR correction* (Benjamini & Hochberg)”, “*Conservative stats*”. Intersections between lists of microRNAs or targets were performed using R software (v.3.4.0).

### 8. Detection of microRNAs by RT-qPCR

We evaluated the following fractions from CML patients or HD samples: CML LSC CD34^+^CD38^-^CD26^+^, CML HSC CD34^+^CD38^-^CD26^-^, CML progenitors CD34^+^CD38^+^, HD HSC CD34^+^CD38^-/dim^, HD progenitors CD34^+^CD38^+^. We applied a modification of the protocol reported by Chen *et al*, based on a reverse transcription (RT) using gene-specific stem-loop primers^49^ (incubation times were modified as detailed below), followed by individual qPCR reactions for each microRNA using an intercalating agent. qPCR used a specific forward primer and a universal reverse primer designed to hybridize with the constant region included in the stem-loop primer. Two multiplex RT reactions for microRNAs (M1 and M2) (see Supplementary Table S1) were performed for each sample. Additionally, each sample was reverse transcribed with random primers in a separate reaction, in order to measure snRNA U6 as a reference gene for qPCR. Final concentrations of components of RT reaction were: dNTPs 0.25mM (Invitrogen or INBIO Highway); DTT 10mM (Invitrogen); Superscript II 2.5 U/µL (Invitrogen); RNAse inhibitor 0.2 U/µL (RNAseOUT, Invitrogen); *stem-loop* primer 0.05 µM (each) or random primers 0.01 µg/µL (Invitrogen). Incubation times were: 5 minutes of RNA, H2O, and dNTPs at 65°C; tubes were immediately placed on ice; the remaining components were added to the tube and incubated for 30 minutes at 16°C, 40 cycles (30 seconds at 30°C + 30 seconds at 42°C + 1 second at 50°C), followed by a final step of 5 minutes at 85°C. cDNA was diluted (1/2) with distilled water and stored at -20°C. 2 µL of diluted cDNA was used for each qPCR reaction, using the following conditions: forward primer 0.3 µM; universal reverse primer 0.3 µM, and SYBR Green (PowerUp SYBR Green MasterMix, Applied Biosystems; according to the information provided by the manufacturer, Mg^2+^ concentration can vary between 4.76-6.44 mM); incubated for 2 minutes at 50°C, 2 minutes at 95°C, 50 cycles (15 seconds 95°C + 1 minute at 60°C), in a Rotor-Gene Q qPCR equipment (Qiagen). Melting curves were evaluated in order to assess specificity of the reaction. Quantifications were performed in duplicate. In cases were duplicate measurements differed (ΔCt>2), a triplicate measurement was performed. RT-qPCR efficiency was estimated by performing curves of RNA; formula used for efficiency estimation was E= [10^(−1/*m*)]-1, *m* being the slope of the curve. Sequences of all primers used for quantification of microRNAs are available (see Supplementary methods).

### 9. Statistical analysis and graphical tools

GraphPad Prism 6, Microsoft Excel 2007, and Inkscape 0.92 software were used for graphics. Infostat v.2018e software (Córdoba, Argentina) was used for statistical analysis. Data from quantification of microRNAs by RT-qPCR were analyzed using the variable dCt= (Ct microRNA X - Ct snRNA U6), with a linear mixed-effects model (ANAVA): fraction (LSC, CML HSC, CML progenitors, HD HSC, HD progenitors) was set as a fixed effect, and sample (each patient or HD) was set as a random effect (correlation factor: compound symmetry). Variance was modelled using “VarIdent” function (using the variable “fraction”). False discovery rate was considered by multiplying p-values by the number of total microRNAs evaluated. *A posteriori* comparisons were performed using DCG formula^50^.

## Availability of data and materials

All data analysed during this study are included in this published article (and its supplementary information files).

## List of abbreviations

aUPD: acquired uniparental disomy
BFU-E: burst-forming unit-erythroid
BM: bone marrow
CFU-GEMM: colony-forming unit-granulocyte erythroid macrophage megakaryocyte
CFU-GM: colony-forming unit-granulocyte macrophage
CML: chronic myeloid leukemia
FACS: fluorescent-activated cell sorting
GO: Gene Ontology
HD: healthy donor
HSC: hematopoietic stem cells
KEGG: Kyoto Encyclopedia of Genes and Genomes
LSC: leukemic stem cells
LT-HSC: (murine) long-term repopulating hematopoietic stem cell
NGS: next-generation sequencing
PB: peripheral blood
MNC: mononuclear cells
Ph: Philadelphia chromosome
Pre-L-NSC: pre-leukemic neoplastic stem cells
qPCR: quantitative PCR
RT: reverse transcription
snoRNAs: small nucleolar RNAs
snRNAs: small nuclear RNAs
TKI: tyrosine kinase inhibitors
tRNAs: transfer RNAs

## Acknowledgements

We thank CML patients, healthy donors, and the service of Hemotherapy from Instituto Alexander Fleming (Buenos Aires, Argentina) for their invaluable help by providing samples for this project, and Gerardo Cueto for his advice on statistical analysis of data. This work was supported by grants from CONICET-FAPERJ (2013), Agencia Nacional de Promoción Científica y Tecnológica of Argentina (PICT-2013 1710), Fundación Mosoteguy, and Fundación SALES. PV and his team are supported by the Austrian Science Fund FWF, grant F4704-B20. M.B., I.L., P.Y. and J.M. are researchers from the Consejo Nacional de Investigaciones Científicas y Tecnológicas of Argentina (CONICET). M.S.R., M.B.S. and D.K. received CONICET fellowships.

## Authors’ contributions

M.S.R. and M.B. designed the experiments. M.S.R. performed most of the experiments and data analysis. M.B.S. contributed to the validation of microRNAs by RT-qPCR and processing of biological samples. S.B. and C.F. contributed to small RNA NGS-library construction and sequencing. D.K. and P.Y. contributed to bioinformatic analysis of NGS-derived data. S.C., M.R.C., J.F., B.M., C.P., M.A.P., A.I.V. and V.V. contributed with patient samples and analysis of clinical data. J.C.S.A., I.Z., I.L. and J.M. participated with helpful discussion and ideas. P.V. contributed conceptual input and technical support in CML stem cell experiments, reviewed the data and drafted parts of the manuscript. M.S.R. wrote the manuscript. M.B. supervised the entire work. All authors read and approved the final manuscript

## Additional information

### Additional material

Supplementary information.docx: supplementary figures, tables and methods

