## Supplementary information for "miRNome profiling of clonal stem cells in Ph^+^ CML"

A1: Centro de investigaciones oncológicas – Fundación Cáncer, Ciudad Autónoma de Buenos Aires, Argentina.
A2: Centro de investigaciones oncológicas – Fundación Cáncer, Ciudad Autónoma de Buenos Aires, Argentina.
B1: Centro de Transplante de Medula Óssea, Instituto Nacional de Câncer, Rio de Janeiro, Brazil.
C: Programa de Genética, Instituto Nacional de Câncer, Rio de Janeiro, Brazil.
D1: Instituto de Investigación en Biomedicina de Buenos Aires (IBioBA)-CONICET-Partner Institute of the Max Planck Society, Ciudad Autónoma de Buenos Aires, Argentina.
D2: Instituto de Investigación en Biomedicina de Buenos Aires (IBioBA)-CONICET-Partner Institute of the Max Planck Society, Ciudad Autónoma de Buenos Aires, Argentina.
E1: Instituto Alexander Fleming, Ciudad Autónoma de Buenos Aires, Argentina.
E2: Instituto Alexander Fleming, Ciudad Autónoma de Buenos Aires, Argentina.
F1: Hospital Nacional Posadas, El Palomar, Buenos Aires, Argentina.
G1: Hospital J. M. Ramos Mejía, Ciudad Autónoma de Buenos Aires, Argentina.
H: Hospital Interzonal General de Agudos, Prof. Dr. R. Rossi, La Plata, Buenos Aires, Argentina.
I: Fundaleu, Ciudad Autónoma de Buenos Aires, Argentina.
G2: Hospital J. M. Ramos Mejía, Ciudad Autónoma de Buenos Aires, Argentina.
F2: Hospital Nacional Posadas, El Palomar, Buenos Aires, Argentina.
E3: Instituto Alexander Fleming, Ciudad Autónoma de Buenos Aires, Argentina.
J: Instituto de Medicina Experimental, CONICET/Academia Nacional de Medicina, Ciudad Autónoma de Buenos Aires, Argentina.
B2: Centro de Transplante de Medula Óssea, Instituto Nacional de Câncer, Rio de Janeiro, Brazil.
A3: Centro de investigaciones oncológicas – Fundación Cáncer, Ciudad Autónoma de Buenos Aires, Argentina.
K: Department of Internal Medicine I, Division of Hematology and Hemostaseology, Medical University of Vienna, Vienna, Austria.
L: Ludwig Boltzmann Institute for Hematology and Oncology, Medical University of Vienna, Vienna, Austria
A4: Centro de investigaciones oncológicas – Fundación Cáncer, Ciudad Autónoma de Buenos Aires, Argentina.

*: corresponding authors. Full postal address: Centro de investigaciones oncológicas – Fundación Cáncer. Conesa 1003/1005, C1426DRB, Ciudad Autónoma de Buenos Aires, Argentina.

**Supplementary figures.**

Sup. Fig. S1. Gating strategy used for LSC and HSC cell sorting from BM or PB CML patients at diagnosis.

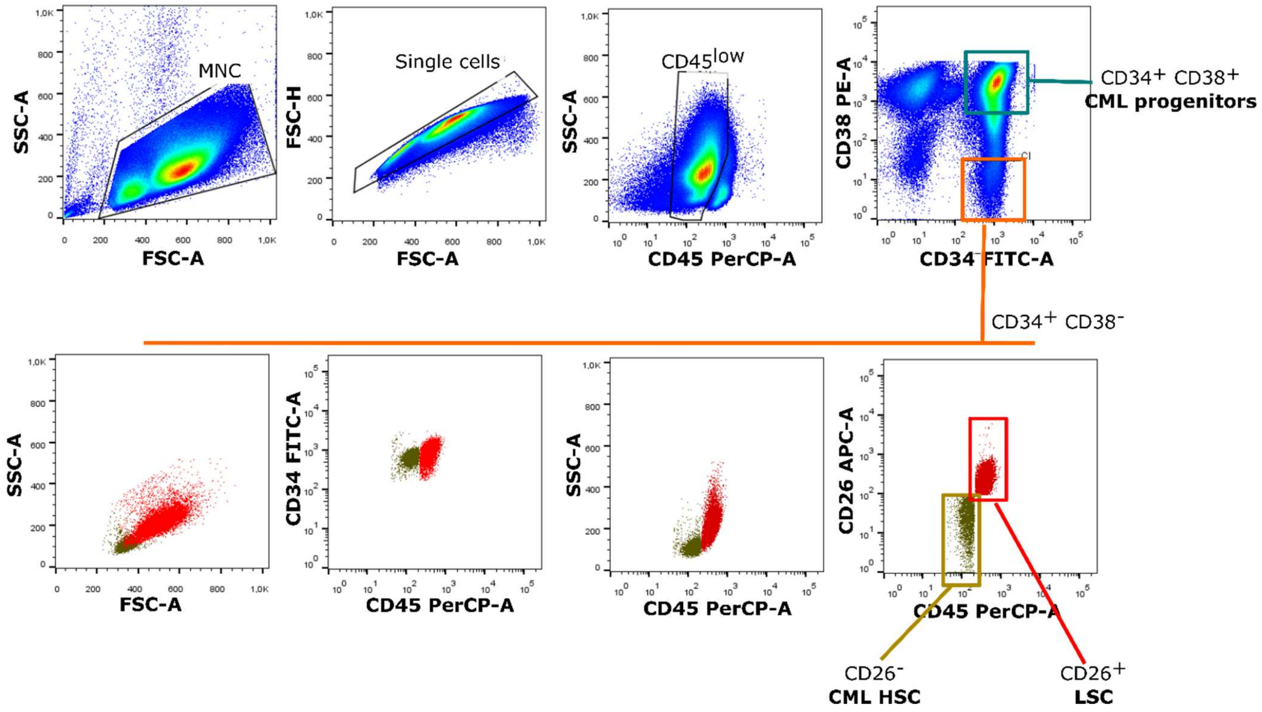

Sup. Fig. S2. Representative flow cytometry patterns observed in $CD34^{+}CD38^{-}$ fractions from CML patients at diagnosis. Pattern 1 showed coexistence of both LSC and HSC populations; pattern 2 showed mostly HSC; pattern 3 showed a single population with varying levels of CD26. Assessment of purity of each fraction revealed that $CD26^{-}$ cells in pattern 3 expressed *BCR-ABL1* mRNA and were not further used for microRNA isolation. Bottom: Each dot represents a pool of 4-6 colonies. CML1, CML2 and CML3 correspond to three single patients with flow cytometry patterns 1, 2 or 3, respectively.

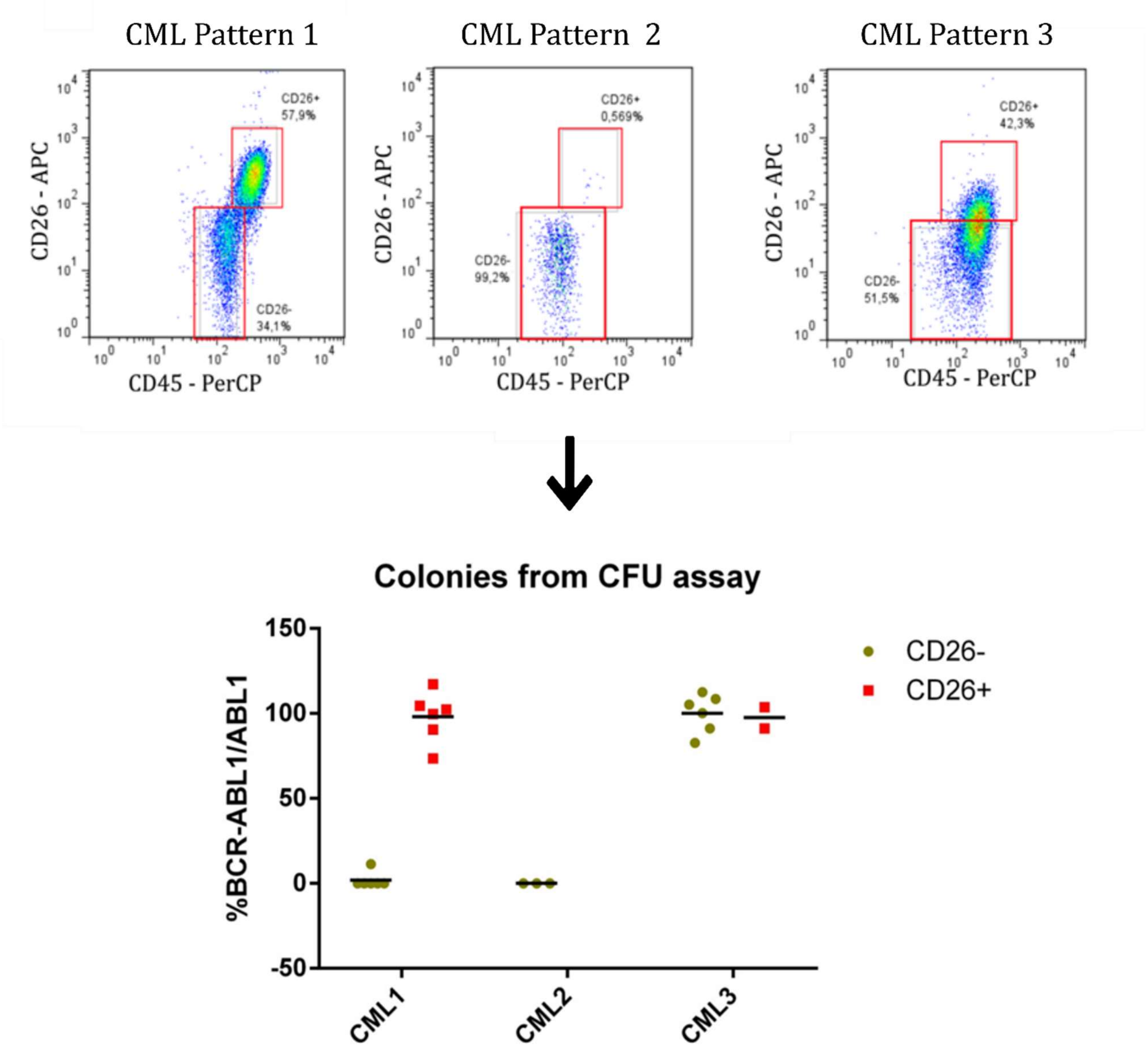

Sup. Fig. S3: Assessment of purity by detection of *BCR-ABL1* mRNA in sorted fractions used for validation (RT-qPCR in a new cohort of samples). Each dot represents a different fraction from all patients evaluated.

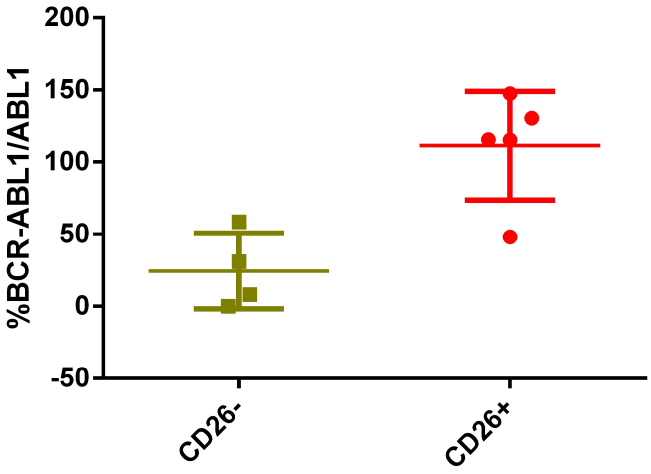


Sup. Fig. S4. Validation of microRNAs by RT-qPCR in a new cohort of CML and HD samples: results for miR-132-3p, let7a-5p, and "novel-3". Results are expressed as ΔCt = Ct (microRNA) - Ct (snRNA U6). Each dot is the mean of technical duplicates from each patient or HD. CML samples are represented in grey symbols, and HD samples in green symbols. Lines connect different fractions from the same patients or HD. n.s.= not statistically significant differences (linear mixed-effects model, *a posteriori* comparison, global α = 0.05)

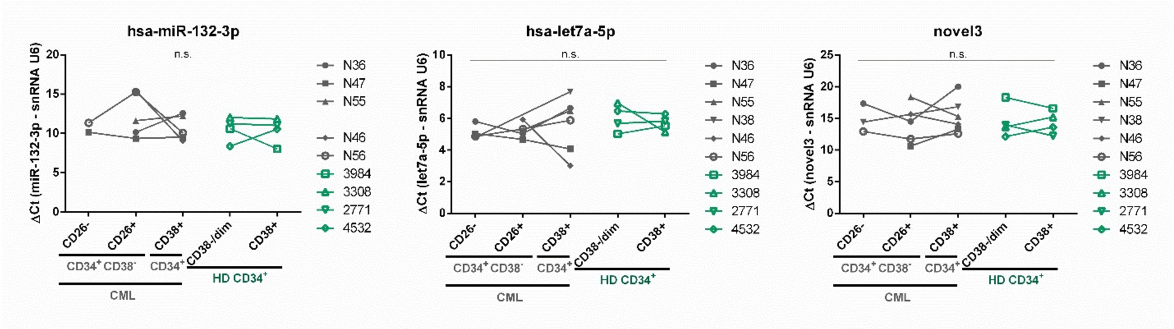

Sup. Fig. S5. Comparison of NGS and RT-qPCR results. Mean $Log_2$(Fold change) was calculated between LSC and CML HSC samples. Dotted line indicates $Log_2$(Fold change) =1 or -1.

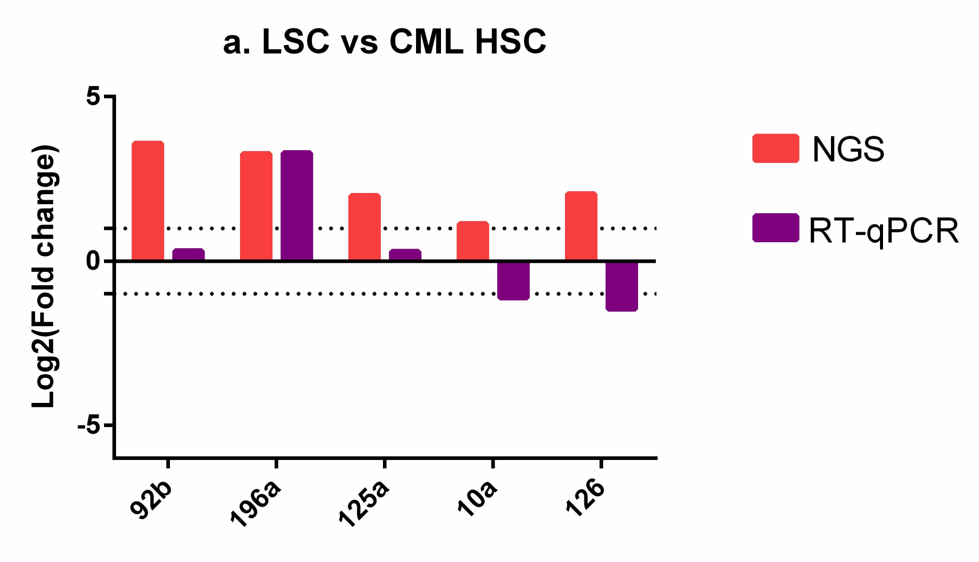


Sup. Fig. S6. Mean $Log_2$(fold change) values for all samples evaluated by RT-qPCR between CML progenitors ($CD38^+$) and LSC (red), CML HSC (blue), HD progenitors (purple) and HD HSC (green), for each microRNA.

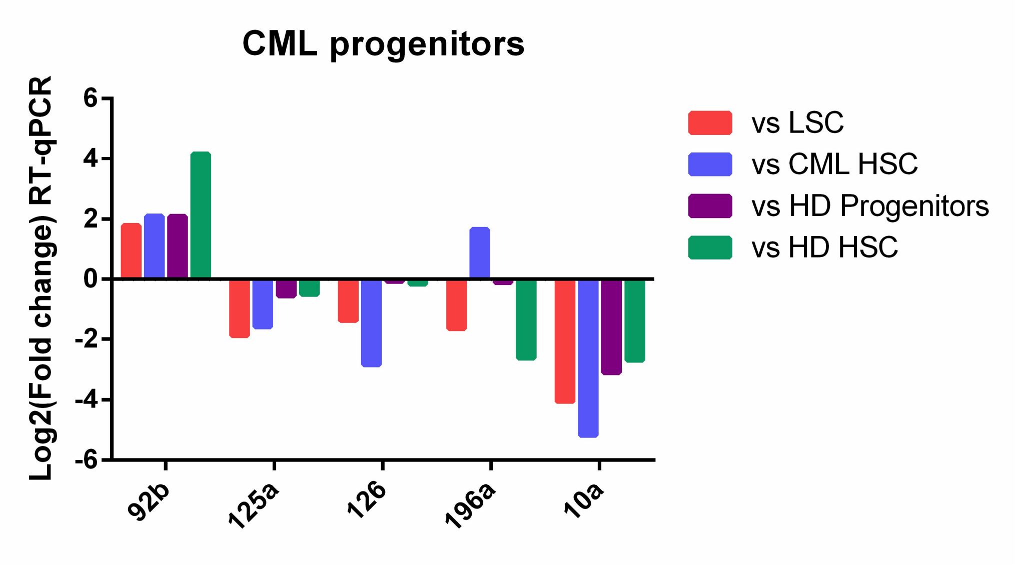

### Supplementary methods

*Red cell lysis of mononuclear cells*

Cells were resuspended in 1mL of pre-warmed (37°C) red cell lysis solution: ethylenediaminetetraacetic acid (EDTA) 0.13mM, potassium bicarbonate ($KHCO_3$) 1mM, ammonium chloride ($NH_4Cl$) 170mM, pH 7.3, and vigorously mixed for 1 minute. After addition of 9mL of red cell lysis solution, cells were incubated 3 minutes at 37°C. Cells were washed by addition of 30mL of phosphate buffered saline (PBS) and centrifuged.

*Isolation of CD34$^+$ cells*

Cells were incubated with 100 µL of beads (CD34 MicroBeads, Miltenyi Biotech), and 100 µL of FcR-block for 30 minutes, for every 1x$10^8$ MNC (in ~300 µL of ice-cold MACS buffer), in a water bath at 2-8°C. All subsequent steps were performed with cold MACS buffer (Bovine Serum Albumin 0.5%, EDTA 2mM, PBS, pH 7.2): cells were washed twice (4 mL), then seeded in a MS column (Miltenyi Biotech) in a volume of 500 µL, washed four times (1x 1 mL, 3 x 500 µL), and eluted in a 2 mL volume. Cells were immediately incubated with antibodies for FACS sorting, or freezed in 1mL of cold freezing medium: Dulbecco's Modified Eagle Medium (DMEM, GIBCO) 50%/ Human Serum Albumin 40%/ Dimethyl sulfoxide 10%. During thawing of CD34$^+$ cells, cells were quickly thawed in a water bath at 37°C, diluted in 30mL of Roswell Park Memorial Institute 1640 medium (RPMI 1640, GIBCO), centrifuged, resuspended in 50-100 µL of MACS buffer, counted and immediately incubated with antibodies for FACS sorting.

*Assessment of purity in sorted fractions.*

Total RNA was extracted from CFU assay-derived colonies following manufacturer's instructions (RNAqueous-Micro Kit, Ambion). RNA was eluted in 18 µL of pre-warmed (75°C) elution solution, and reverse transcribed using Superscript II enzyme (Invitrogen) and random primers (final concentration: 20 ng/µL, Invitrogen), in a reaction volume of 25 µL, following manufacturer's instructions. cDNA was diluted (1/2) with distilled water, and kept at -20°C. Detection of *BCR-ABL1* and *ABL1* was performed by qPCR using primers (final concentration: 300 nM each, Invitrogen) and probes (final concentration: 200nM, SIGMA-Aldrich) recommended by the "Europe against cancer" program, Taqman Universal MasterMix II, no UNG (Applied Biosystems), by means of an Applied Biosystems 7500 Real-time PCR machine.  Absolute copy numbers and amplification efficiency were estimated by the inclusion of calibration curves built with standardized plasmids (BCR-ABL pDNA calibrant; ERM certified reference material ERMAD623, SIGMA).

*Primers used for quantification of microRNAs by RT-qPCR*

| | |
|---|---|
| Stem-loop primers for reverse transcription of microRNAs | |
| hsa-miR-92b-3p RT | GTCGTATCCAGTGCAGGGTCCGAGGTATTCGCACTGGATACGACGGAGGC |
| hsa-miR-125a-5p RT | GTCGTATCCAGTGCAGGGTCCGAGGTATTCGCACTGGATACGACTCACAG |
| hsa-miR-182-5p RT | GTCGTATCCAGTGCAGGGTCCGAGGTATTCGCACTGGATACGACAGTGTG |
| novel-3 RT | GTCGTATCCAGTGCAGGGTCCGAGGTATTCGCACTGGATACGACCAAGAC |
| hsa-miR-2355-5p RT | GTCGTATCCAGTGCAGGGTCCGAGGTATTCGCACTGGATACGACTTGTCC |
| hsa-miR-126-5p RT | GTCGTATCCAGTGCAGGGTCCGAGGTATTCGCACTGGATACGACCGCGTA |
| hsa-miR-132-3p RT | GTCGTATCCAGTGCAGGGTCCGAGGTATTCGCACTGGATACGACCGACCA |
| hsa-let-7a-5p RT | GTCGTATCCAGTGCAGGGTCCGAGGTATTCGCACTGGATACGACTTCTAT |
| hsa-miR-10a-5p RT | GTCGTATCCAGTGCAGGGTCCGAGGTATTCGCACTGGATACGACCACAAA |
| hsa-miR-196a-5p RT | GTCGTATCCAGTGCAGGGTCCGAGGTATTCGCACTGGATACGACCCCAAC |
| Forward primers for qPCR | |
| hsa-miR-92b-3p Fw | TGCATTTCTATTGCACTCGTC |
| hsa-miR-125a-5p Fw | TGATCCCTGAGACCCTTTAAC |
| hsa-miR-182-5p Fw | GGTTTGGCAATGGTAGAACT |
| novel-3 Fw | CGGACTGAAGGTAGATAGAACAG |
| hsa-miR-2355-5p Fw | TCGAAGATCCCCAGATACAAT |
| hsa-miR-126-5p Fw | CAGCGGCATTATTACTTTTGG |
| hsa-miR-132-3p Fw | CTCGGTAACAGTCTACAGCCA |
| hsa-let-7a-5p Fw | GCGGTGAGGTAGTAGGTTGT |
| hsa-miR-10a-5p Fw | GGCTACCCTGTAGATCCGAA |
| hsa-miR-196a-5p Fw | GCGTCGTAGGTAGTTTCATGTT |
| snRNA U6 Fw | GCTTCGGCAGCACATATACTAAAAT |
| Reverse primers for qPCR | |
| Universal Rv (microRNAs) | GTGCAGGGTCCGAGGT |
| snRNA U6 Rv | CGCTTCACGAATTTGCGTGTCAT |

### Supplementary tables

Sup. Table S1. Experimental design of multiplex RT of microRNAs performed during validation by RT-qPCR.

| Multiplex 1 (M1) | Multiplex 2 (M2) |
|---|---|
| miR-92b-3p<br>miR-125a-5p<br>miR-182-5p<br>novel-3<br>miR-2355-5p<br>miR-126-5p | miR-132-3p<br>let-7a-5p<br>miR-10a-5p<br>miR-196a-5p |
| Positive control: KU812 | Positive control: TF1a |

RT of microRNAs was performed on two multiplex reactions (M1 and M2), each including different combinations of stem-loop primers. KU812 and TF1a cell lines were used as positive controls.

Sup. Table S2. MicroRNAs dysregulated in LSC by small RNA-NGS.

| | GFOLD value | | DESeq normalised counts | | |
|---|---|---|---|---|---|
| | **CML HSC vs LSC** | **HD HSC vs LSC** | **LSC** | **HSC (CML)** | **HSC (HD)** |
| hsa-mir-92b-3p | -3.6 | 1.6 | 2062 | 207 | 5951 |
| hsa-mir-196a-5p | -3.6 | -2.7 | 135 | 6 | 13 |
| hsa-mir-126-5p | -2.0 | 1.1 | 18667 | 5343 | 44262 |
| hsa-mir-125a-5p | -1.9 | -1.0 | 6225 | 2114 | 3045 |
| hsa-mir-2355-5p | -1.5 | -1.0 | 928 | 385 | 444 |
| hsa-mir-99b-5p | -1.2 | -1.7 | 3907 | 2214 | 1207 |
| hsa-mir-411-5p | -1.2 | 3.6 | 36 | 9 | 749 |
| hsa-mir-10a-5p | -1.1 | 1.1 | 28770 | 19727 | 68583 |
| hsa-mir-708-5p | 1.6 | 2.5 | 4 | 46 | 42 |
| hsa-mir-431-5p | 1.6 | 1.9 | 8 | 112 | 89 |
| hsa-mir-134-5p | 1.9 | 2.5 | 1 | 64 | 62 |
| hsa-mir-485-3p | 2.2 | 2.7 | 1 | 53 | 46 |
| hsa-mir-409-3p | 2.6 | 3.8 | 46 | 595 | 902 |
| hsa-mir-323b-3p | 2.8 | 2.2 | 3 | 140 | 60 |
| hsa-mir-432-5p | 2.9 | 2.4 | 0 | 105 | 65 |
| hsa-mir-382-5p | 3.7 | 2.9 | 3 | 130 | 52 |

GFOLD can be considered as a robust $Log_2$(fold change) value. GFOLD<0: increased in LSC; GFOLD>0: decreased in LSC.

Sup. Table S3. Molecular pathways associated to targets of selected microRNAs with detectable counts and GFOLD=0 between LSC and CML HSC.

| List 1: hsa-mir-3136, hsa-mir-543, hsa-mir-3202, hsa-mir-1256, hsa-mir-1976, hsa-mir-1185-1-3p, hsa-mir-639, hsa-mir-550a-3p, hsa-mir-548b-5p, hsa-mir-3136-5p, hsa-mir-939-5p, hsa-mir-3134 hsa-mir-375, hsa-mir-1908-5p, hsa-mir-202-5p, hsa-mir-885-5p | | | |
|---|---|---|---|
| KEGG pathway | p-value | #genes | #microRNAs |
| Fatty acid biosynthesis | 7.77E-16 | 1 | 1 |
| List 2: hsa-mir-612, hsa-mir-650, hsa-mir-632, hsa-mir-564, hsa-mir-1225-5p, hsa-mir-135a-5p, hsa-mir-3140-3p, hsa-mir-3200-3p. hsa-mir-1909-5p, hsa-mir-760, hsa-mir-3144-3p, hsa-mir-3125, hsa-mir-3161, hsa-mir-3121-3p, hsa-mir-611, hsa-mir-1252-5p | | | |
| KEGG pathway | p-value | #genes | #microRNAs |
| Extracellular matrix-receptor interaction | 0 | 22 | 3 |
| Hippo signaling pathway | 2.21E-01 | 7 | 2 |

.

Sup. Table S4. snoRNAs dysregulated in LSC by small RNA-NGS.

| | GFOLD value | |
|---|---|---|
| | **HD HSC vs LSC** | **HD HSC vs. CML HSC** |
| 14q(II-24) | 0.6 | 1.1 |
| 14q(II-21) | 1.0 | 0 |
| 14q(II-15) | 1.1 | 1 |
| 14q(II-22) | 3.4 | 2.5 |
| 14q(II-9) | 3.5 | 2.2 |

GFOLD can be considered as a robust $Log_2$(fold change) value. HD HSC vs LSC: GFOLD<0= increased in LSC, GFOLD>0= decreased in LSC. HD HSC vs CML HSC: GFOLD<0= increased in CML HSC, GFOLD>0= decreased in CML HSC

Sup. Table S5. MicroRNAs dysregulated in CML HSC by small RNA-NGS.

| | GFOLD | DESeq normalised counts | |
|---|---|---|---|
| | **HD HSC vs CML HSC** | **HSC (CML)** | **HSC (HD)** |
| hsa-mir-1304-3p | -3.6 | 1179 | 765 |
| hsa-mir-629-5p | -2.3 | 2841 | 421 |
| hsa-mir-629-3p | -2.3 | 1431 | 167 |
| hsa-mir-362-5p | -2.0 | 7186 | 1204 |
| hsa-mir-20a-5p | 2.0 | 4054 | 16670 |
| hsa-mir-195-5p | 2.1 | 16 | 104 |
| hsa-mir-654-3p | 2.4 | 67 | 516 |
| hsa-mir-224-5p | 2.5 | 2 | 36 |
| hsa-mir-144-3p | 2.6 | 415 | 3244 |
| hsa-mir-181c-5p | 2.7 | 2315 | 12831 |
| hsa-let-7c-5p | 2.7 | 236 | 896 |
| hsa-let-7f-5p | 2.9 | 23477 | 139037 |
| hsa-mir-548o-3p | 3.0 | 11 | 157 |
| hsa-let-7a-5p | 3.0 | 17636 | 120313 |
| hsa-mir-182-5p | 3.5 | 4567 | 41803 |
| hsa-mir-183-5p | 3.8 | 328 | 4196 |

GFOLD can be considered as a robust $Log_2$(fold change) value. GFOLD<0: increased in CML HSC; GFOLD>0: decreased in CML HSC.
